# Phosphate overplus response in *Chlamydomonas reinhardtii*: polyphosphate dynamics to monitor phosphate uptake and turnover

**DOI:** 10.1101/2024.01.25.577272

**Authors:** Tatiana Zúñiga-Burgos, Adolfo Saiardi, Miller Alonso Camargo-Valero, Alison Baker

## Abstract

Many micro-organisms store inorganic phosphate (Pi) in the form of polyphosphate (polyP) and exhibit in-cell polyP accumulation, a phenomenon known as ‘phosphate overplus response’, when resupplied with Pi after a period of deprivation. Quantitative and qualitative methods were used to follow the dynamics of polyP synthesis and turnover in four strains of *Chlamydomonas reinhardtii* during Pi deprivation followed by nutrient resupply. The lowest level of in-cell polyP during Pi deprivation, which also correlates with the cessation of growth, is the key parameter for the timing of Pi resupply to maximise the Pi overplus response Additional nutrients do not affect the size of the overplus response, but they are important for continued growth and maximal Pi removal from the media. Tracking polyP allows the correct time for nutrient resupply to be determined and therefore a reproducible Pi overplus response to be achieved. Depending on whether maximum cellular phosphorus (P) content or maximum Pi removal is desired different strategies may be required – e,g., Pi deprivation until growth cessation then resupplying complete nutrients gives the best trade-off between high in-cell P accumulation, high Pi uptake and algal biomass growth. Although polyP levels are maintained after Pi resupply, the polymer is dynamically remodelled. IP_6_ increases during this time. This increase does not precede polyP synthesis as predicted by a model where inositol phosphates switch on polyP synthesis. One strain tested, CC-5325, shows enhanced Pi uptake and levels of polyP and total in-cell P, suggesting that strain selection is also important.

**Importance:** There is strong interest in using microalgae to sustainably control and recover nutrients, especially P, from wastewater. This would help to meet environmental discharge consents and recycle nutrients into agriculture or other applications. Like bacteria and yeasts, microalgae exhibit a Pi overplus phenomenon when Pi-deprived cells are resupplied with P, but microalgae do not require an additional carbon source and can simultaneously uptake nitrogen as well. Use of microalgae in wastewater treatment is limited by the unpredictability of their response and sensitivity to environmental factors, but engineered systems can greatly benefit from better understanding Pi dynamics and polyP accumulation. In the literature there is a lack of consensus regarding protocols to maximise the Pi overplus. In this work we provide robust measurements of quantitative physiological parameters, which should allow reproducibility in laboratory studies and provide design parameters for algal-based nutrient recovery systems from waste waters.

## Introduction

Microalgae have evolved to survive in environments where nutrient supply fluctuates in space and time. As a result, they have complex homeostatic mechanisms that enable them to integrate nutrient acquisition, storage and growth which are still incompletely understood (1–4). Inorganic phosphate (PO_4_^3-^, there after abbreviated as Pi) is an essential molecule, required for nucleic acids, phospholipids, energy metabolism and cell signalling. It is acquired from the environment by phosphate transporters and then utilised for biosynthesis of macromolecules. In microorganisms it is stored predominantly in the form of polyphosphate (polyP). This capacity for storage allows survival under conditions of phosphate scarcity (5).

Microalgae, like bacteria and yeasts, exhibit a ‘Pi overplus response’ when they are transferred to a plentiful supply of phosphate following a period of phosphate deprivation (6, 7). Recent studies in the model alga *Chlamydomonas reinhardtii* are shedding light on some of the mechanisms by which phosphate is sensed and transported and polyP is synthesised. PolyP is an orthophosphate polymer widespread in nature (8–11). The synthesis of polyP by *C. reinhardtii* is conjectured to be like *Saccharomyces cerevsiae* and occurs through the VTC complex in the vacuolar membrane (12, 13). In *S.cerevisiae*, phosphate transported into the cell drives ATP synthesis, which in response triggers production of inositol phosphates (IPs). IPs bind to the SPX domain of the VTC complex to activate polyP synthesis (14–16). However, *C. reinhardtii* differs from yeast regarding polyP turnover, since the exopolyphosphatase Ppx1 has no homologues in microalgae (17, 18). The mechanisms of Pi sensing and response are also distinct, with the myb transcription factor Phosphate Starvation Response 1 (PSR1) (19, 20), which is homologous to vascular plant PHR1 being a key mediator of phosphate starvation responses in algae, instead of the PhoR/PhoB regulon in *E.coli* and Pho80/Pho85/Pho4 in *S.cerevisiae* (21). Other than a phosphate reserve, polyP is considered a source of energy (22) and a cation chelator assisting in the resistance to heavy metals (23, 24). PolyP may also act as a regulator of enzyme activity and adaptation to stress responses and late-growth phases (25, 26).

The ability of microorganisms to hyperaccumulate phosphate as polyP has been exploited for removal of nutrients from wastewater. Enhanced Biological Phosphate Removal (EBPR) at wastewater treatment works relies on the activity of polyphosphate accumulating organisms (PAOs) under a combination of aerobic and anaerobic conditions – i.e., in an anaerobic phase, PAOs take up energy-rich volatile fatty acids (e,g., acetate) and store them as polymers, such as polyhydroxyalkanoates, while under aerobic conditions, they degrade glycogen to drive uptake of phosphate and accumulation as polyP (27). The EPBR process requires a source of carbon (acetate), whereas microalgae are capable of photoautotrophic or mixotrophic growth. Thus, there is a lot of interest in harnessing the ability of microalgae to take up and store phosphate as a means of reducing the carbon footprint of wastewater treatment (28–30). However, the performance of microalgae in this regard is unpredictable due to the lack of solid underpinning knowledge regarding the physiological responses of microalgae and the allocation of Pi between different pools under conditions of nutrient deprivation and resupply. Most published studies carry out nutrient deprivation for a specified time, without measuring any specific physiological parameter (7, 31–34). This makes comparing the results practically impossible.

To address this limitation, quantitative and qualitative tools for polyP analysis in *Chlamydomonas* were developed and used to characterise and quantify in-cell polyP and relate these changes to standardised measures of growth, other in cell Pi pools and nutrient uptake. This revealed the dynamics of polyP synthesis and turnover, and the role of nutrients other than Pi in the overplus response and allowed definition of conditions to maximise the Pi overplus response in *C. reinhardtii*.

## Results

### Inorganic phosphate reserves (polyP) are used before organic phosphate pools (RNA) after resuspension in Pi-free media

To study Pi deprivation in *C. reinhardtii,* we monitored biomass concentration, cell counts, RNA and polyP in four different strains to determine the time when cessation of growth and lowest internal Pi occur following resuspension in Pi-free media (TA) (**Figure 1A-D** and **Table 1**.). For all the four strains, growth continues for 24 h after resuspension, independently of Pi availability. The strain CC-4350, which has much higher cell numbers than the other strains, has cells that are typically small meaning that it requires a higher number of cells to generate the same biomass (Chlamydomonas Resource Center - http://chlamycollection.org). We applied a quantitative method to determine polyP based on digestion of an RNA preparation extracted from *Chlamydomonas* cells with recombinant *S.cerevisiae* Ppx1 polyphosphatase (see materials and methods). All strains rapidly depleted polyP reserves within the first 24 h of Pi deprivation (**Figure 1F)**. The lowest polyP content is maintained at approximately 0.10% PO_4_-P, on dry weight (dw) basis, except for the strain CC-4350 (0.06% PO_4_-P, dw). Conversely, after resuspension in TAP, cells maintained a constant polyP content, which was higher in strain CC-5325 (**Figure 1E**). In cells resuspended in TA media (**Figure 1H**), RNA levels declined as cells entered stationary phase and did not fall below 2.08% RNA, dw, even after 96 h of Pi deprivation. A similar decline in RNA content in the biomass is observed in the absence of Pi deprivation (**Figure 1G**), although the overall RNA content in the biomass was higher (Significance between resuspension in TA or TAP, 24 and 96 h time points, *p* value = 3.60×10^-8^ and 1.43×10^-7^, respectively).

**Figure 1.**
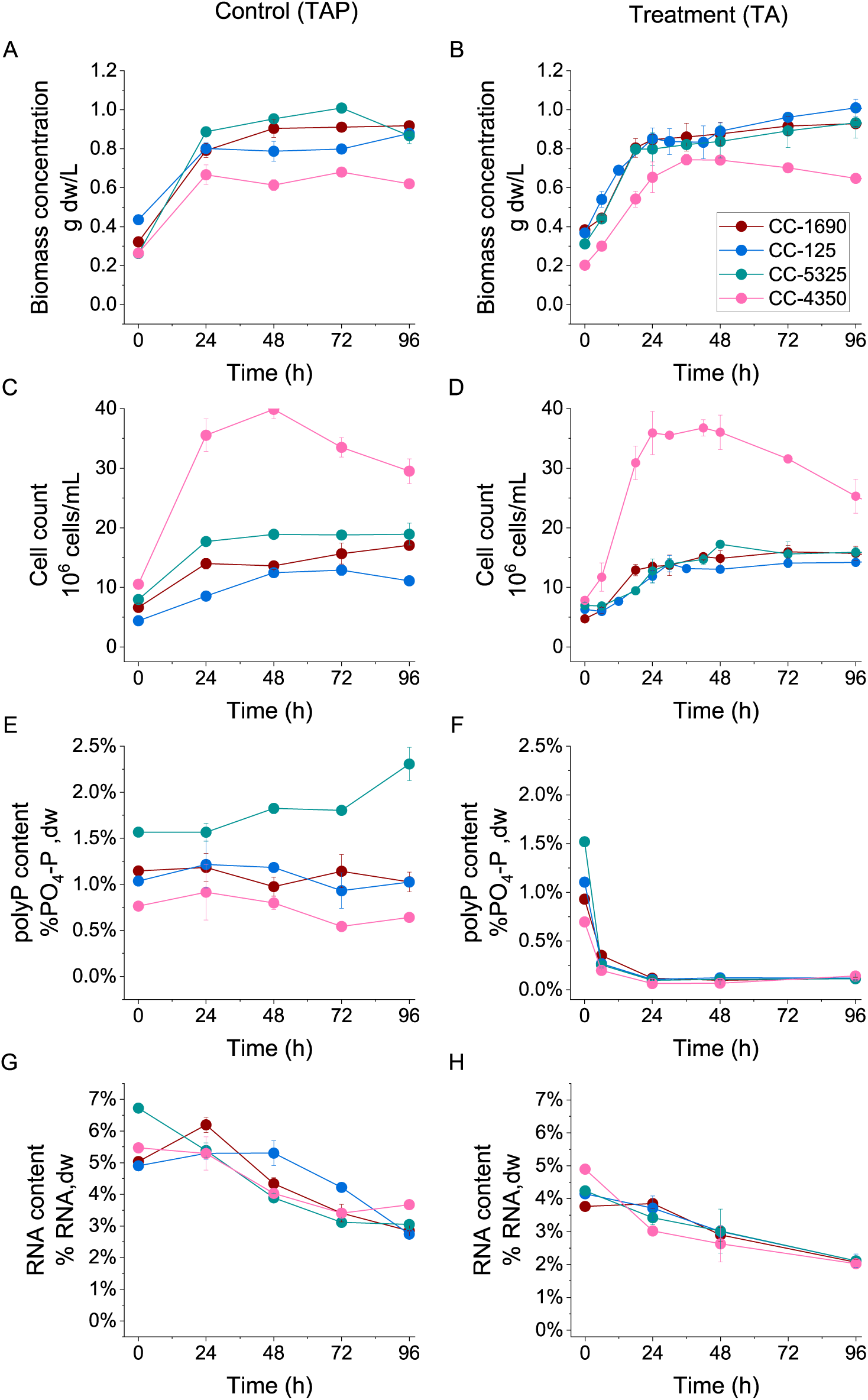
Phosphate deprivation results in rapid decrease of polyP content in *Chlamydomonas* cells. Cells grown in TAP media until OD_750nm_= 1.0, harvested (washed) and resuspended in TAP (left) or TA (right). A and B, Biomass concentration (g dw/L), C and D, Cell counts (10^6^ cells/mL), E and F, polyP content in the biomass (% PO_4_-P, dw), and G and H, RNA content in the biomass (% RNA,dw)

**Table 1.** Algal biomass growth characteristics during Pi deprivation. OD_750nm_= 1.0 cells grown in TAP media were harvested (washed) and resuspended in TAP or TA media.

| A |  |  |
| --- | --- | --- |
| Specific growth rate (h<sup>-1</sup>) |  |  |
| Strain | Control (TAP) | Treatment (TA) |
| CC-1690 | 0.037 ± 0.002 | 0.032 ± 0.002 |
| CC-125 | 0.027 ± 0.002 | 0.032 ± 0.003 |
| CC-5325 | 0.051 ± 0.001 | 0.041 ± 0.002 |
| CC-4350 | 0.038 ± 0.003 | 0.044 ± 0.004 |
| Average | 0.038 ± 0.005 | 0.037 ± 0.003 |
| B |  |  |
| Doubling time (h) |  |  |
| Strain | Control (TAP) | Treatment (TA) |
| CC-1690 | 24.2 ± 5.0 | 20.1 ± 3.1 |
| CC-125 | 25.2 ± 2.8 | 24.1 ± 2.2 |
| CC-5325 | 20.9 ± 2.7 | 25.9 ± 2.2 |
| CC-4350 | 13.8 ± 1.2 | 11.0 ± 1.2 |
| Average | 21.0 ± 2.6 | 20.3 ± 3.3 |
| C |  |  |
| Biomass productivity (g dw L<sup>-1</sup> h<sup>-1</sup>) |  |  |
| Strain | Control (TAP) | Treatment (TA) |
| CC-1690 | 0.021 ± 0.002 | 0.019 ± 0.001 |
| CC-125 | 0.018 ± 0.001 | 0.020 ± 0.002 |
| CC-5325 | 0.027 ± 0.001 | 0.020 ± 0.002 |
| CC-4350 | 0.018 ± 0.002 | 0.019 ± 0.002 |
| Average | 0.021 ± 0.002 | 0.020 ± <0.001 |

Total P content in the biomass (**Figures 2A** and **S2A**), shows the same trend as polyP. All the strains reach a minimum total P content of ∼0.50% PO_4_-P,dw, after 24 h of resuspension in TA media (**Figure S2A**). Presumably, this decline is due initially to polyP and later RNA degradation, although polyP is not completely depleted from the cells. Whereas the polyP:total P ratio is maintained at ∼50% when cells have access to Pi in the media (**Figure 2B**), the decline in polyP levels during Pi deprivation decreases the polyP:total P ratio to a minimum of ∼20% after 6 h for the strains CC-5325 and CC-4350, and after 24 h for the CC-1690 and CC-125 strains (**Figure S2B)**. Initial Pi concentration in the media (31.9 ± 0.5, mg P/L) decreased during the first 48 h after resuspension in TAP (**Figure 2C**). During this period, the strains CC-1690, CC-125 and CC-4350, removed 44% of Pi on average from the media, whereas the strain CC-5325 removed an average of 87% of initial PO_4_-P, which is consistent with their higher total P content in the biomass (approx. 3.5% P, dw) compared to the other strains (approx. 2.0% P, dw) (**Figure 2A**).

**Figure 2.**
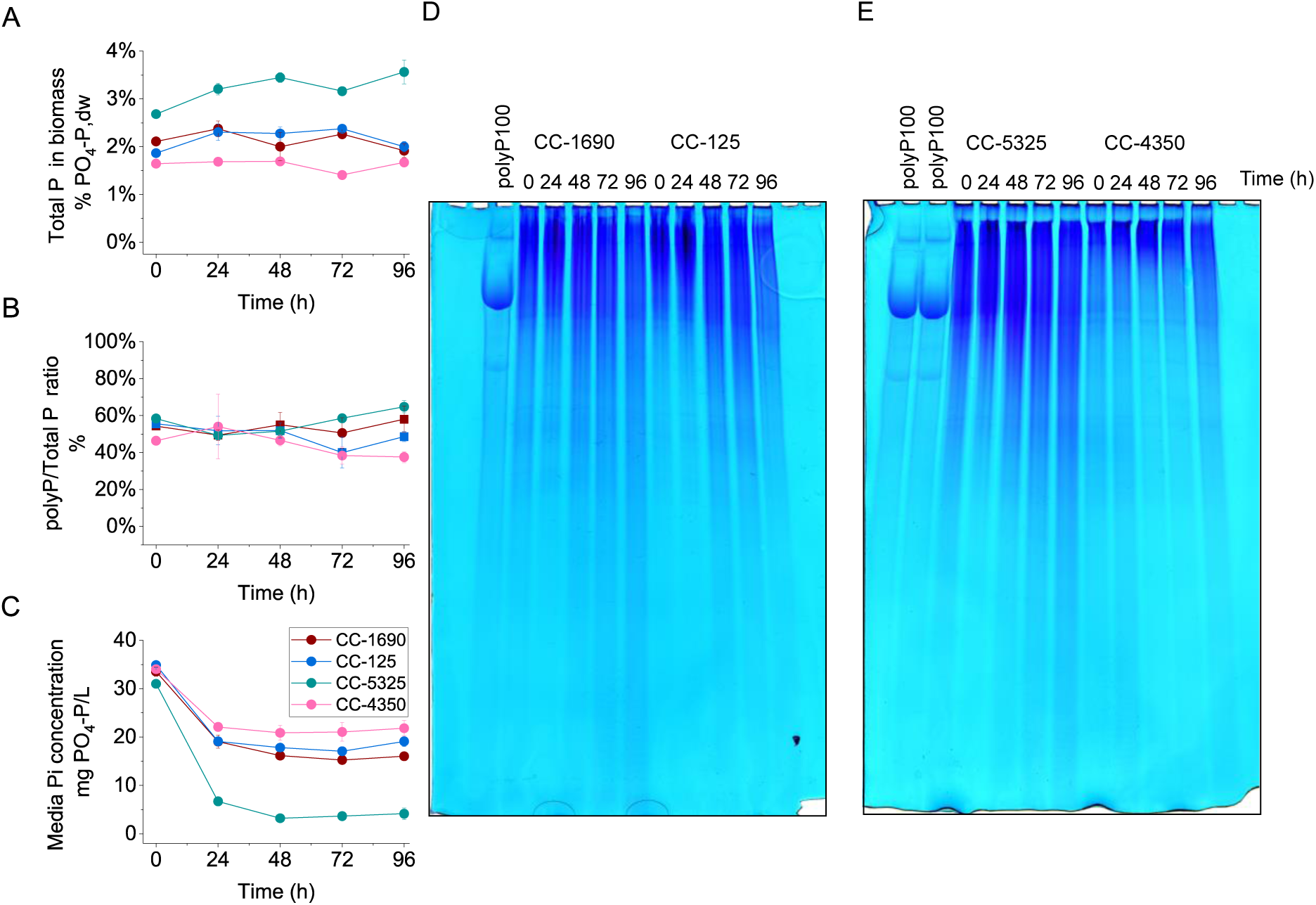
Under Pi-replete conditions strains vary in their ability to take up Pi and store as polyphosphate. A Total P in biomass (% PO_4_-P,dw), B Mass ratio of polyP in proportion to total P in biomass (%), C. Phosphate concentration in the media (mg PO_4_-P/L). D 20% PAGE. Left, polyP100 loading control, for the four strains of *Chlamydomonas* the RNA volume equivalent to 500µg dw was loaded on each well for the 0, 24, 48, 72 and 96 h time points after OD1.0 cells were harvested and resuspended in TAP media.

Polyacrylamide gel electrophoresis allows a qualitative view of polyphosphate as a polymer, which complements the quantitative assay. Long chain polyP runs at the top of the gel, while the smear represents medium or low molecular weight polyP. **Figure S2C** shows a consistent decline in polyP after resuspension in Pi free media. This is in contrast with **Figure 2D**, where no decrease in polyP is observed when cells were resuspended in fresh TAP media, although long polyP chains are degraded into medium-to short-chain polyP, after biomass growth ceased. In accordance with the polyP quantification shown in **Figure 1E**, the strains CC-5325 and CC-4350 contain the highest and the lowest polyP concentration, respectively.

### Only Pi repletion is required to trigger Pi overplus, but complete nutrients are required for enhanced biomass production

Cessation of growth and the lowest polyP content occurs after 24h of Pi deprivation in all strains. At this point cells were supplied with 1 mM Pi, either by addition of KPO_4_ solution or harvested and resuspended in fresh TAP media. **Figure 3A** shows biomass growth during the first 12 h after repletion with all nutrients (TAP media), which led to an average biomass productivity of 0.026 g dw L^-1^ h^-1^ for all the strains (**Table S1**). The average biomass productivity in the control experiment (no Pi deprivation) was 0.021 g dw L^-1^ h^-1^ (**Table 1**). No difference was observed in the specific growth rates of the four strains when resuspended in TAP (**Table S1**). Biomass growth was negligible when Pi was supplied as KPO_4_ (Biomass productivity <0.005 g dw L^-1^ h^-1^) consistent with Pi not being limiting for growth (**Table 1**). The RNA content in the biomass shows an initial small recovery followed by a decline as the cultures enter stationary phase (**Figure 3B**) The cultures resuspended in TAP had a significantly higher RNA content in the biomass than cultures repleted with KPO_4_, 24 h after repletion (*p* value=0.03) but there was no difference by the end of the experiment (96 h; *p* value =0.81).

**Figure 3.**
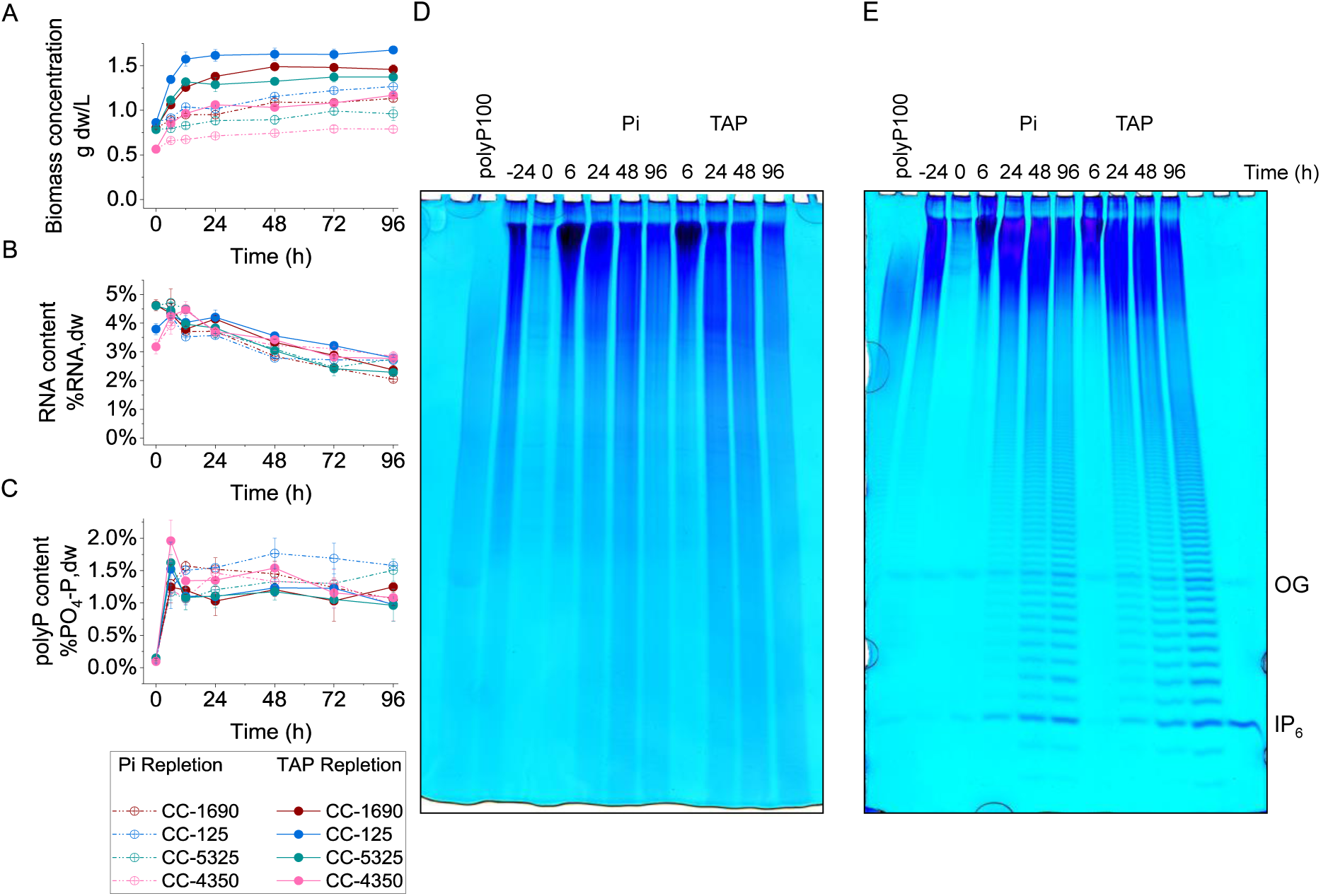
PolyP reserves are rapidly resynthesized on provision of Pi after 24 h of Pi deprivation. OD_750nm_= 1.0 cells were harvested (washed) and resuspended in TA media. After 24 h, the cultures were repleted to 1mM Pi with either KPO_4_ solution (dashed dotted lines) or resuspended in fresh TAP media (solid lines), and monitored for 96 h. A Biomass concentration (g dw/L), B RNA content in biomass (% RNA, dw), C. PolyP content in the biomass (% PO_4_-P,dw). D. 20% (left) and 33% (right) representative PAGE of CC-1690 RNA volume equivalent to 500 µg dw. Wells from left to right: polyP100 loading control, -24 h (OD1.0 cells before Pi deprivation), 0 h (24 h Pi-deprived cells before Pi repletion), 6-96 h (After repletion with KPO_4_), 6-96 h (After resuspension in TAP media) and 2 nmol of IP_6_ standard. IP_6_ overplus can be observed by increasing PAGE concentration to 33% (right). OG; Orange G marker.

All four strains exhibited a 15-fold increase in their polyP content after only 6 h following Pi resupply (**Figure 3C**), and there was no difference between repletion type (*p* value =0.13). Total P content in the biomass, shows a 6-fold increase in this same period. In contrast to the polyP response, cultures repleted with TAP had significantly higher total P content (*p* value =0.001) than those repleted with KPO_4_, at the 6 h time point (**Figure S3A**). The polyP content, compared with total in-cell P was restored to the ∼50% benchmark (see **Figure S3B** and compare with **Figure 2B**). Analysis of polyP by PAGE for the strain CC-1690 (**Figure 3D)** shows a rapid polyP synthesis in the first 6 h of the repletion phase followed by turnover (**Figure 3D**). The pattern in Pi overplus observed in these gels is consistent with the other three strains CC-125, CC-5325 and CC-4350 (**Figure S4**).

### Longer Pi deprivation altered RNA content in the biomass and triggered lower polyP accumulation upon repletion

We next analysed total RNA content in the biomass and observed that Pi deprivation for 96 h led to a ∼50% reduction in the RNA content of cells (**Figure 1H**). Repletion with TAP after 96 h of Pi deprivation resulted in biomass growth for all the strains, but growth was negligible when cultures were resupplied with Pi alone (Biomass productivity <0.005 g L-1 h-1) (**Figure 4A**), like responses of cells Pi-deprived for 24 h (**Figure 3A**). The change in biomass RNA content after repletion at 96 h (**Figure 4B**) is completely different. Repletion with TAP leads to RNA recovery that matches the biomass growth period (between 12-24 h after Pi supply). However, after the maximum value is reached at 24 h, RNA levels in the biomass fall, and by 72 h they reached the level before Pi repletion. CC-4350 appears to be an outlier but is affected by the low biomass concentration compared to the other strains. In contrast, in cultures repleted with Pi only, RNA levels do not recover. While a Pi overplus response is seen in cells that had been Pi-deprived for 96 h, it is only 6-fold (**Figure 4C**), compared to the 15-fold increase seen with cells that had been Pi-deprived for only 24 h (**Figure 3C**). This response was observed in all strains, independently of whether Pi only or TAP was supplied. Pi resupply to cells that had been Pi-deprived for 96 h led to less accumulation of total P compared to cells that had only been deprived of Pi for 24 h (**Figure S3C**), and consequently a lower polyP:total P ratio (**Figure S3D**). Analysis of samples from strain CC-1690 by PAGE showed resynthesis of polyP followed by remobilisation (**Figure 4D**), like the samples from 24 h Pi-deprived cells (**Figure 3D**) but with lower intensity consistent with the quantitative data. The other strains behaved similarly to CC-1690 (**Figure S5**).

**Figure 4.**
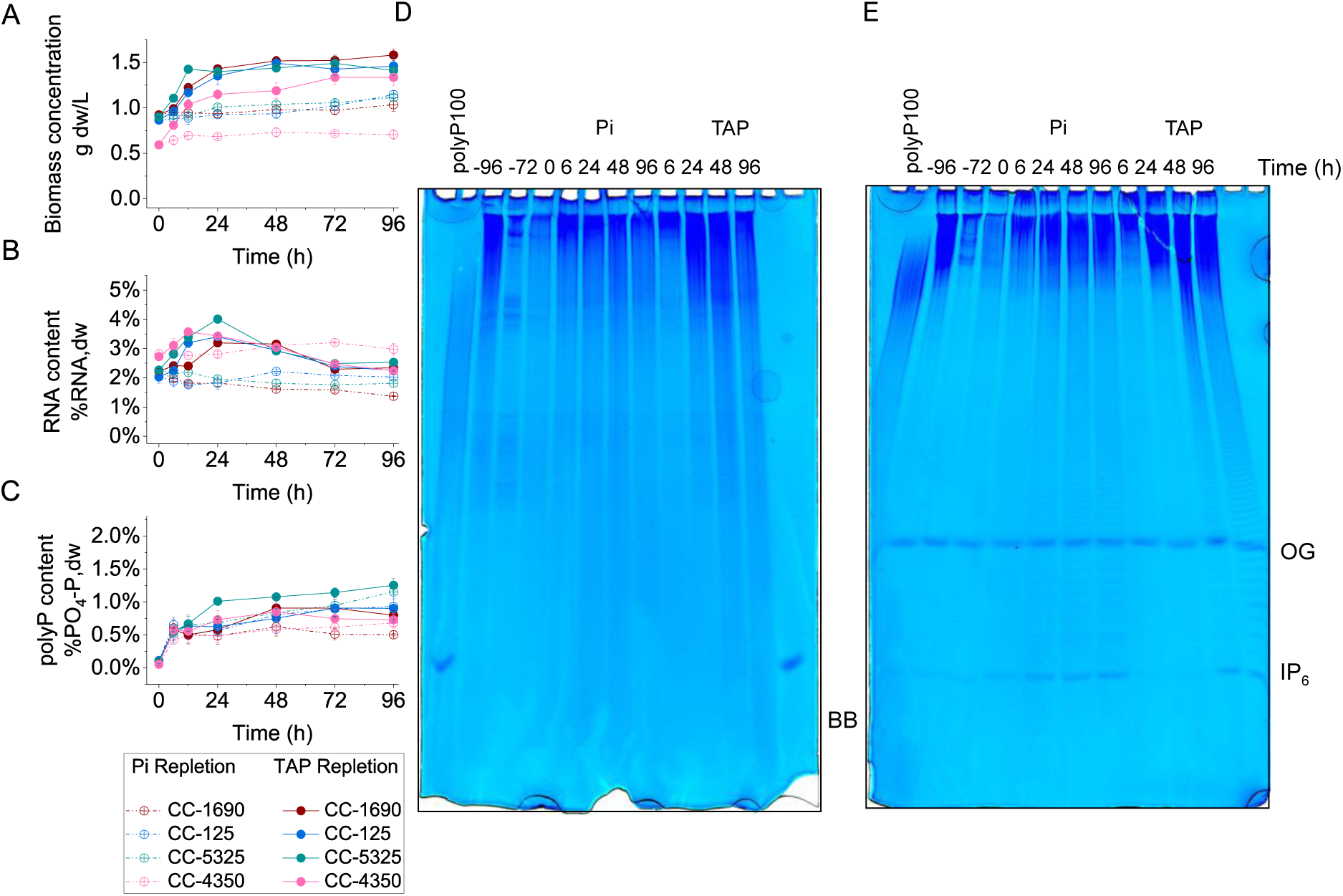
Longer Pi deprivation led to a lower Pi overplus independent of Pi repletion type but RNA resynthesis required complete nutrients. OD_750nm_= 1.0 cells were harvested (washed) and resuspended in TA media. After 96 h, the cultures were repleted to 1mM Pi with either KPO_4_ solution (dashed dotted lines) or resuspended in fresh TAP media (solid lines), and monitored for 96 h. A Biomass concentration (g dw/L), B RNA content in biomass (% RNA, dw), C. PolyP content in the biomass (% PO_4_-P,dw). D. 20% (left) and 33% (right) representative PAGE CC-1690 RNA volume equivalent to 500 µg dw. Wells from left to right: polyP100 loading control, -96 h (OD1.0 cells before Pi deprivation), -72 h (After 24 h Pi deprivation), 0 h (96 h Pi-deprived cells before Pi repletion), 6-96 h (After repletion with KPO_4_), 6-96 h (After resuspension in TAP media) and 2 nmol of IP_6_ standard. IP_6_ overplus can be observed by increasing PAGE concentration to 33% (right). BB bromophenol blue marker. OG Orange G marker.

### Inositol hexakisphosphate (IP_6_) does not precede polyP accumulation

A faint IP_6_ signal could be observed on 33% PAGE 6 h after repletion with both KPO_4_ and TAP at both 24 h (**Figure 3D**) and 96 h (**Figure 4D**) time points. The intensity of the IP_6_ signal increases with time and reaches its maximum at the end of the monitored period. This pattern is the same in all four *C. reinhardtii* strains **(Figures S4** and **S5**) and is independent of phosphate repletion type. The antibiotic neomycin is an inhibitor of phospholipase C which generates IP_3_, the precursor for IP_6_ and has been used to inhibit production of IPs in *C. reinhardtii* (6, 35). Since previous work that used neomycin failed to test the effect on IP_6_ synthesis, we decided to monitor the effect of the antibiotic on IP_6_ production (**Figure 5**). IP_6_ level was unaffected by neomycin, but cell growth was inhibited with an effect visible after 6 h of neomycin treatment.

**Figure 5.**
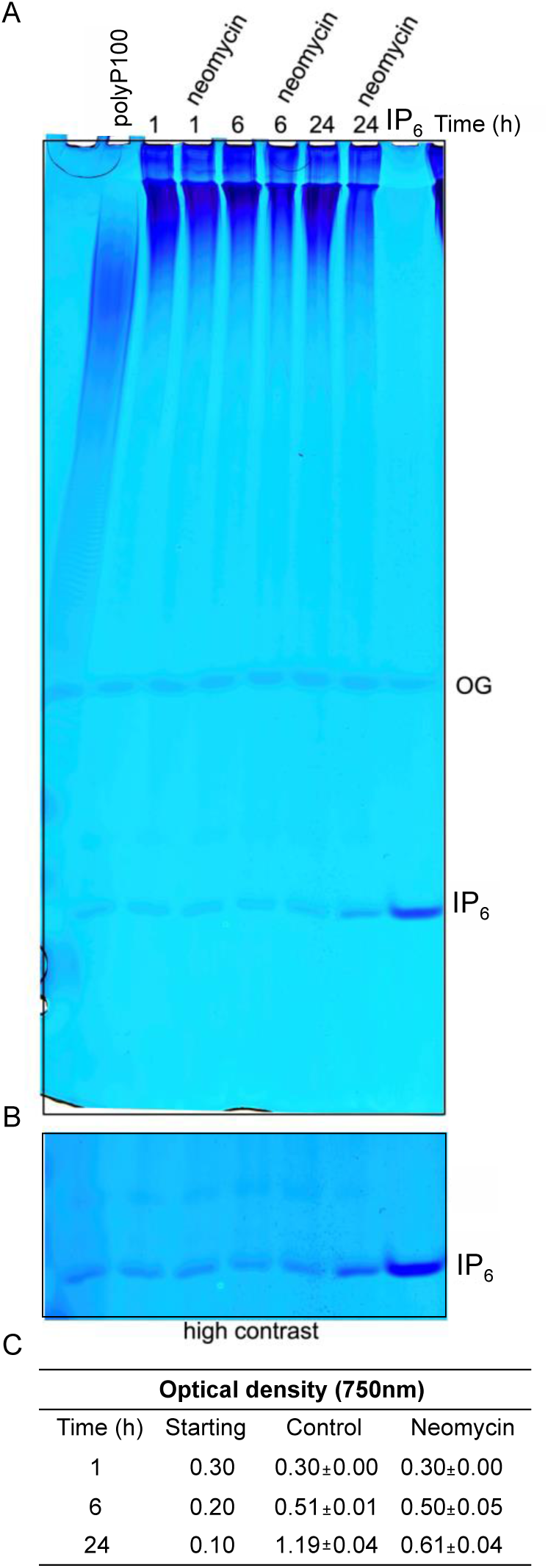
Neomycin treatment does not inhibit IP_6_ synthesis but is toxic to *Chlamydomonas* cells. Fresh *C. reinhardtii* CC-5325 (OD_750nm_= 0.6-0.8) culture were diluted into 30 ml in fresh TAP media. To take into account cell division, a differential initial inoculation was used of OD_750nm_ 0.3, 0.2 and 0.1 for 1, 6 or 24 hours treatment respectively. Neomycin was used at 20 µM. A 33% PAGE gel with 20 µg RNA loading of samples collected after 1, 6 and 24 h of 20 µM Neomycin treatment, against a control, 2 nmol IP_6_ was use as running control. B High contrast of the PAGE shown above for improve visualisation of IP_6_ bands. C. OD_750nm_ measured during the time course experiment shows exponential growth of control, in contrast with growth cessation, 24 h after Neomycin treatment.

### Enhanced Pi removal from the media occurs upon Pi refeeding with all other nutrients

The removal of Pi from the media was monitored throughout the experiment. Supply of TAP (solid lines) following 24 h Pi deprivation led to rapid and complete removal of phosphate from the media, whereas supplying Pi alone resulted in a lower removal (approx. 62% average Pi removed for the strains CC-5325, CC-125 and CC-4350 and 38% for CC-1690) (**Figure 6A**).

**Figure 6.**
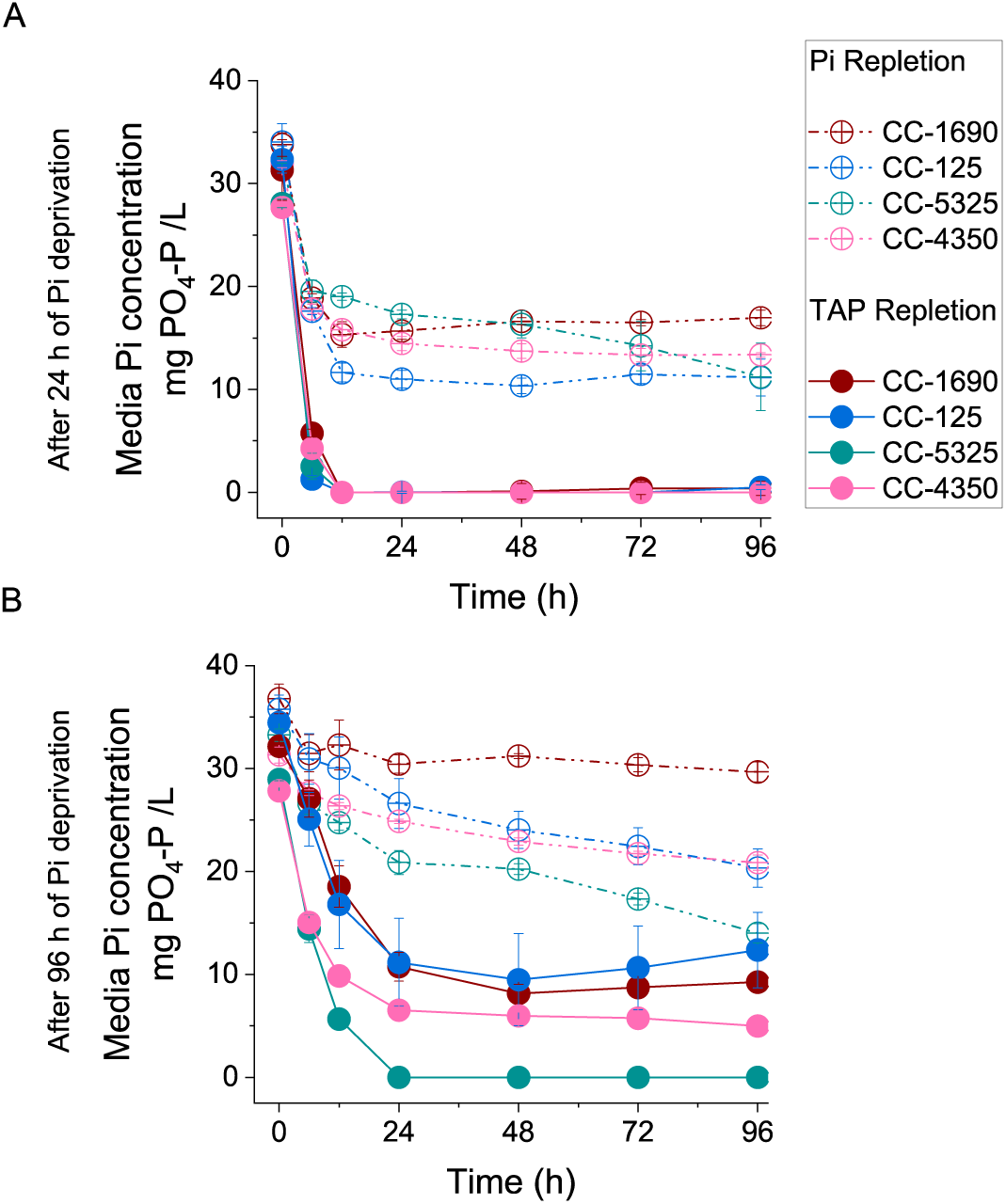
Provision of complete nutrients is required for effective phosphate uptake following Pi deprivation. Phosphate removal from the media (mg PO_4_-P/L) in Pi-repleted cells after A 24 h of Pi deprivation and B 96 h of Pi deprivation. OD1.0 cells were harvested (washed) and resuspended in TA media. Pi-deprived cells were repleted with 1 mM Pi with either KPO_4_ (dashed dotted lines) or resuspended in fresh TAP media (solid lines).

Cells which had been Pi-deprived for 96 h removed Pi more slowly and less extensively than cells that had been Pi-deprived for 24h, but the difference between Pi only and TAP repletion was retained for all strains. (**Figure 6B**). The strain CC-5325 was the only strain able to remove all Pi from the media following a 96 h Pi deprivation treatment when resupplied with TAP, and did this within 24 h. Overall, enhanced Pi removal coincides with biomass growth when Pi repletion occurred as resuspension in fresh TAP media.

To further assess the effect of the addition of other nutrients apart from Pi on the nutrient uptake rates, a correlation-based principal component analysis (PCA) was carried out. This tool helped to reduce the dimensionality across both periods of Pi deprivation and Pi repletion types. PCA allows the overall correlation between the nutrient uptake rates of phosphate (as P source), ammonium (as N source), and sulphate (as S source) with their initial concentrations in the media to be established (**Figure 7**). The nutrient uptake rates were calculated in segments (0-6 h, 6-12 h and 12-24 h) after Pi repletion (**Table S2**) using the media concentration data from **Figure 6** and **S7**. PC1 and PC2 in **Figure 7** accounted for 91.9% of the variance across the data. PC1 (representing 71.5% of the correlation), explains the effect of the addition of N and S in the nutrient uptake rates. The loading vectors of initial N and S concentration in the media are positively correlated with PC1 and with all three nutrient uptake rates. PC2 (representing 18.4% of the correlation) accounts for the effect of Pi addition only, and only Pi uptake rate has a positive correlation with this principal component, whereas N and S uptake rates are negligible to PC2. The principal component scores (dots) represent the Pi repletion time segments (0-6 h, 6-12 h and 12-24 h) for 24 and 96 h Pi-deprived *Chlamydomonas* cultures, repleted with Pi only or TAP. The scores are consistently allocated according to the availability of N and S (PC1) or the addition of Pi (PC2). The strongest correlations were observed during the first six hours of Pi repletion, when the most rapid changes were observed. As time passed (6-12 h and 12-24 h), the strength of correlation to each component weakens. It is interesting, given that for both periods of Pi deprivation, polyP accumulation was no different if Pi was supplied together with all other nutrients (TAP media) (**Figure 3C** and **4C**). Pi uptake rate is positively correlated with both PC1 and PC2. However, when Pi-deprived cells were supplied with all nutrients in TAP, more Pi was taken up together with other nutrients supplied to sustain biomass growth (**Figures 3A** and **4A**).

**Figure 7.**
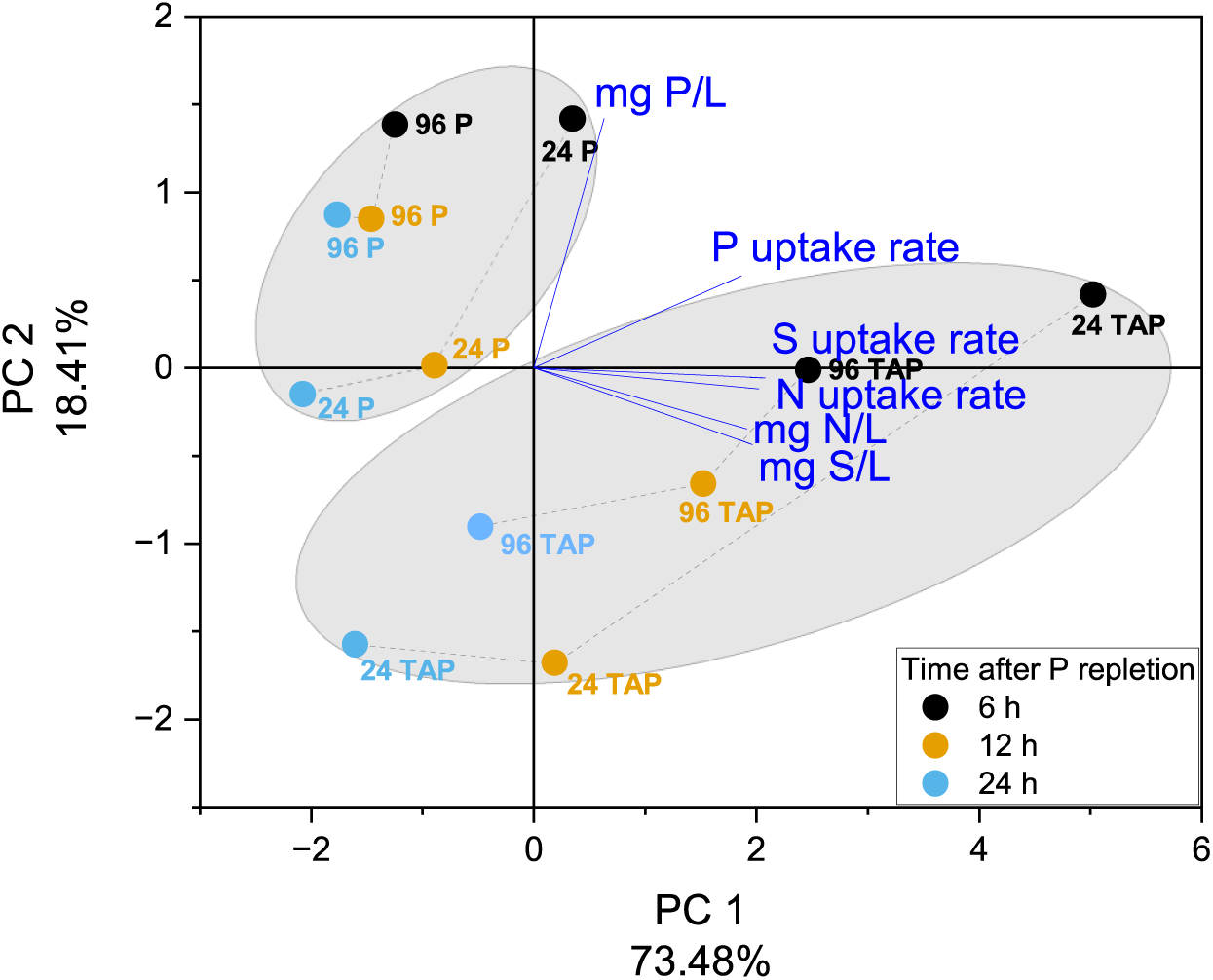
Correlation-based PCA analysis is shown for the average of four strains of *C. reinhardtii* (CC-1690, CC-125, CC-5325 and CC-4350). Biplot shows the principal component 1 (x-axis) explaining 73.48% and principal component 2 (y-axis) explaining 18.41% of the correlation of P, N and S uptake rates with the concentration of nutrients in the media (PO_4_-P, NH_4_-N, SO_4_-S in mg/L). The loading vectors close together indicate the parameters with more similar variance. The dots show the values calculated for each of the six parameters (loading vectors), 6, 12 and 24 h after repletion of 1 mM Pi with either Pi (KPO_4_ solution) or resuspended in fresh TAP media, of 24 or 96 h Pi-deprived cells. Ellipses group the dots and loading vectors according to Pi repletion type.

## Discussion

This study explores the relationship between the duration of Pi deprivation, the turnover of polyP and RNA, the timing of repletion of Pi and the effect of repleting with Pi only or with complete media (TAP). Several conclusions can be drawn from the data. With respect to the timing of Pi repletion, the maximum overplus response was achieved by resupplying nutrients at the point of lowest polyP content, which also coincided with the cessation of growth. In our experiments this was 24 h following Pi removal and the size of the Pi overplus (15-fold) was independent of the type of repletion (Pi only or TAP) and independent of the strain tested. The Pi overplus response was entirely due to resynthesis of polyP which occurred within just 6 h of Pi resupply. Growth cessation was not due to exhaustion of nutrients (including phosphate) in the media in non-Pi deprived cells (**Table 1**) in accordance with previous reports for *Chlorella*. In *Chlorella* synthesis of polyP is a feature of stationary phase cells resupplied with phosphate and is correlated with upregulation of a putative polyphosphate synthetase (7, 36). Our results show that the internal reserve utilised to sustain growth under Pi depletion conditions is polyP. The ability to specifically quantify polyP in *Chlamydomonas* for the first time also demonstrated that polyP is not completely depleted from the cells, highlighting its other roles apart from P storage (22, 23, 26, 37, 38).

In contrast to polyP, cells initially maintained their RNA levels which declined once stationary phase was reached, independent of whether the cultures were subjected to Pi deprivation or not (**Figure 1G, H**). Presumably once in stationary phase less active metabolism is required. Under Pi-replete conditions cells maintained a polyP:total P ratio of about 50% (w/w) which dropped to about 20% (w/w) after 24 h of Pi deprivation. RNA levels continued to decline over the course of the experiment. When Pi was resupplied at 96 h a Pi overplus response was observed but this was much lower (6-fold) than the response observed when Pi was resupplied at 24 h, the point of minimum polyP and growth cessation. Here also only Pi was required for the overplus response, but RNA levels did not recover. Complete nutrients were required to restore RNA levels and to resume growth. Some studies have reported that a longer Pi deprivation period is required for maximum Pi overplus response (34, 36) but when time is used as the variable it is difficult to compare experiments performed under different experimental conditions. A significant contribution of this study is to demonstrate that lowest in cell polyP corresponds to the point of growth cessation and is optimal time for Pi repletion to maximise the Pi overplus response.

By utilising polyacrylamide gel electrophoresis, the dynamics of polyP and IP_6_ could be observed. The intensity of staining of polyP correlated well with the quantitative measurements, giving confidence of their robustness. PAGE showed that polyP was initially synthesised as high molecular weight polyP and, although polyP levels were maintained when growth ceased after nutrient resupply (presumably cells entered stationary phase due to light limitation as nutrients remained plentiful), polyP was turned over or remodelled into medium and short species. The role of the polyP dynamics and the functions of different polyP chain lengths in *Chlamydomonas* is a question that requires further investigation. IP_6_ was also observed to increase over time after refeeding Pi to Pi-deprived cells. In *S. cerevisiae* IPs bind to the SPX domain of the VTC complex proteins and stimulates their activity as a polyP synthetase (12, 39). The pattern of accumulation of IP_6_ observed here sheds doubt on such a role in *Chlamydomonas*, or at least points to other roles of IP_6_. To try to examine the linkage between polyP synthesis and IP_6_ we used neomycin to attempt to inhibit IPs synthesis as previously described (6). We failed to observe an effect of neomycin on IP_6_ levels, likely due to the presence in Chlamydomonas of the PLC independent cytosolic pathway of IP_6_ synthesis which is dependent on ITPK1 (Inositol-tetrakisphosphate 1-kinase) (40), a potential homologue of which is present in *C. reinhardtii* (NCBI Reference Sequences: XP_042928346). Instead, neomycin was toxic to *Chlamydomonas* at the concentrations used, as visible bleaching of the cells occurred after 24 h of treatment, presumably due to inhibition of chloroplast protein synthesis. To shed further light on this question more specific genetic and pharmacological approaches are needed.

Uptake of Pi from the media is driven by biomass growth and this is stimulated by repletion of all nutrients not just Pi. Depriving cells of Pi until lowest polyP / growth cessation followed by resupply of Pi with complete nutrients led to resumed growth and complete removal of media Pi within 12 h (**Figure 6A**) much more rapid than without prior Pi deprivation (**Figure 2C**) or with longer Pi deprivation (**Figure 6B**). While the effect of Pi deprivation on both Pi overplus and Pi uptake in *Chlamydomonas* has been studied previously and the correlation between increased biomass and Pi uptake noted (34), the role of other nutrients has not been explored despite clear evidence that the responses to Pi, sulphate and nitrogen are interrelated (1, 2, 14, 41, 42). We used PCA to link the correlation of nutrient uptake rates to the initial nutrient concentration of PO_4_-P, NH_4_-N and SO_4_-S according to both Pi deprivation periods and Pi repletion types tested (**Figure 7**). Nutrient uptake rates of N and S, correlate nicely with their own initial concentration in the media as expected. In contrast Pi uptake rate is correlated positively with both N and S, but also Pi concentration in the media. The covariance scores are separated based on nutrient availability, meaning that the effect of nutrient addition clearly explained most of the nutrient removal after Pi repletion. The covariance scores show that the most rapid changes occurred during the first 6 h after Pi supply as KPO_4_ or TAP. This allowed TAP repleted cultures to produce biomass, hence removing more Pi from the media, than cultures repleted with KPO_4_.

In almost all respects there was no difference in behaviours of the four *Chlamydomonas* strains tested. CC-1690 and CC-125 are standard widely used laboratory strains with a normal cell wall. CC-5325 and CC-4350 are cell wall deficient (though that has been questioned for CC-5325) (43). CC-5325 is the background strain for the Clip mutant library (44) and it did show one very interesting difference to the other three strains in that it has enhanced Pi uptake and maintains a higher in-cell P level (**Figure 2A** and **C**). The reason for this is unknown but CC-5325 has a mutation in a putative histone deacetylase (44). This could mean that some component involved in Pi sensing or uptake is altered in expression. However, the maximum total P in biomass accumulated by this strain (∼3.5%) is only about 50% of that reported in the strongest over expression line of PSR1 (4). Nevertheless, it demonstrates that strains with higher Pi uptake rates and biomass P levels could be selected for.

Overall, our findings show that mid-exponential phase cultures reached the minimum internal polyP reserve by 24 h after Pi removal. During this period cell division continued and cultures reached early stationary growth. This period of Pi deprivation, together with repletion of Pi together with all other nutrients, achieved the biggest Pi overplus and enhanced Pi removal from the media. From the application perspective, it is more cost effective to use the shortest Pi deprivation period that triggers the best Pi removal. These results suggest that Pi repletion as resuspension in wastewater containing Pi and all other nutrients would be preferable than chemical addition of phosphate. The observation that polyP peaked at 6 h whereas complete Pi removal (24 h Pi deprivation and TAP repletion) was achieved after 12 h also provides information on the potential design parameters. For instance, on when to harvest P-rich biomass and when to bypass Pi-deprived biomass to a new cycle of nutrient removal. These findings require further research to reach the robustness and reproducibility required for real life application. The applicability to other algal strains that might be better suited to WWTW and to situations where a more complex microbial ecology exists as well as to continuous culture systems is also necessary. Nevertheless, our simplified experimental system has allowed quantitative measurement of relevant parameters and dynamics of Pi overplus by microalgae as a phenomenon with the potential to enhance P recovery from wastewater, to contribute to close the phosphorus cycle.

## Materials and Methods

### *Chlamydomonas* sp. strains and cultivation conditions

CC-1690 (also known as 21gr, mt+), CC-125, (mt+ [137c]), CC-5325 (also known as CMJ030, identical to CC-4533 cw15, mt−, CLiP library background (45)) and the arginine auxotrophic strain CC-4350 (cw15 *nit1 nit2 arg7-8* mt+ [Matagne 302]) were obtained from the Chlamydomonas Resource Center (http://chlamycollection.org). For maintenance all strains were cultured axenically in solid Tris Acetate Phosphate (TAP) media without Na_2_SeO_3_ (46), at 25°C, 100 μmol photons m^-2^ s^-1^ constant light. TAP media contained arginine (100 μg/ml) when required. Liquid cultures were shaken at 160rpm, and in all experiments Arginine (100 μg/ml) was added to keep conditions in the culture equivalent for all strains.

### Experimental design

In all experiments, strains were inoculated from a 3-4 day starter TAP liquid culture to an initial OD_750nm_ of 0.005 in 0.75 L TAP media and grown to OD_750nm_ 1.0. To study the effect of Pi deprivation (**Figure S1A**), cultures at OD750nm 1.0 were centrifuged 3000g for 10 min, the cell pellet washed twice with phosphate free media Tris-Acetate (TA) supplemented with KCl to maintain potassium concentration. The biomass was finally resuspended in TA media (200 mL) and transferred to 250 mL conical flasks in triplicates for each strain, to start Pi deprivation. In an otherwise identical control experiment the cell pellet was resuspended in fresh TAP media (**Figure S1B**).

In phosphate overplus experiments Pi deprivation was carried out as described above for either 24 h or 96 h. At the end of Pi deprivation period, cultures were repleted with Pi to trigger P overplus and monitored for 96 h. Each replicate was divided in two equal volumes. One was transferred to a new 250 mL conical flask and supplied with a 1 M solution of potassium phosphate (KPO_4_) composed of 10.8 g K_2_HPO_4_/ 5.6 g KH_2_PO_4_ (w/w, in 100 mL of ddH_2_O) to 1 mM Pi. The second was harvested by centrifugation at 3000g for 10 min, the supernatant was discarded, and the biomass was resuspended in fresh TAP media (1 mM Pi) and transferred to a new 250 mL conical flask with a foam plug (**Figure S1C**). Samples were collected according to **Figure S1D**. Samples (1-40 mL) were centrifuged at 4000g for 5 min and the supernatants sterile filtered (0.22 μm pore size) and stored at -20°C for medium analyses. Biomass pellets were washed twice in ddH_2_O, flash frozen with liquid nitrogen and stored at -70°C.

### Biomass growth and phosphate analyses

OD_750nm_ was monitored at each sampling point. Cell counts determined using a Neubauer chamber. Samples were diluted 10x-20x and fixed using a 10% formaldehyde/0.5 glutaraldehyde solution (v/v). Biomass concentration was obtained by drying biomass pellets with a SpeedVac Plus (SC210a – Thermo Savant Instruments) overnight, and dry weight was determined. A second drying period (overnight) ensured that dry weight data was accurate.

Phosphate analyses in the media and biomass (total P) were performed as described in (4).

### Polyphosphate quantification and detection

RNA was extracted from biomass fresh pellets using phenol-chloroform according to (15), quantified by spectrophotometry (A 260 nm) and kept at -20°C. Quantification of polyphosphate as free phosphate was achieved after enzymatic digestion of 2 μg of RNA with 100 ng recombinant exopolyphosphatase (Ppx1) (15). Soluble phosphate was determined using the malachite green method (47). Untreated RNA and Ppx1 treated samples were diluted 100x with sterile ddH_2_O and loaded in a 96-well plate in technical triplicates, together with a KH_2_PO_4_ calibration curve (0-100 μM Pi). Absorbance was measured with band width of 590-620 nm using a POLARstar OPTIMA plate reader (BMG Labtech).

Polyphosphate concentration was calculated from the calibration curve subtracting any phosphate detected in undigested RNA samples1. Polyphosphate was normalised by dry mass and expressed as %PO_4_-P,dw.

20% and 33.3% Polyacrylamide (PAGE) gels were used to resolve polyP following (48). The RNA volume equivalent to 500 μg biomass dw was loaded together with a polyP100 marker l and a 2 nmol Inositol phosphate 6 standard.

### Data processing and Statistical analysis

The specific growth rates (μ = d^-1^) were calculated during the period where logarithmic growth fitted a linear regression (4). This was 24 h for the Pi deprivation and control experiment and 12 h after Pi repletion. The specific growth rate was determined using the equation μ=(ln(y_1_/y_0_))/(t_1_-t_0_), where y1 and y0 correspond to the biomass concentration values at the beginning (t0) and at the end (t1) of the exponential growth phase. Biomass productivity Bp (g L^-1^ h^-1^) represents the quantity of microalgal biomass generated during the exponential phase. Bp was calculated with the equation Bp=(y_1_-y_0_)/t_1_. Nutrient uptake rates (k=mg g dw^-1^ h) were calculated for Pi overplus experiments for the 0-6 h, 6-12 h and 12-24 h time segments after Pi repletion, as k=(CN_0_-CN_1_)/(y_1_-y_0_)/t_1_-t_0_, where CN_0_ and CN_1_ correspond to the initial nutrient concentration in the media and at the end of the specific period (0-6 h, 6-12 h and 12-24 h).

For the multivariate principal component analysis (PCA), the nutrient uptake rates and the initial nutrient concentration in the media for both Pi deprivation periods, and Pi repletion types tested, were used. The data was normalised by calculating the z-scores by subtracting the value to its mean and dividing it by the standard deviation. The PCA report indicated that 2 principal components contributed to 91.9% of the variance of the data, and eigenvalues for PC selection were higher than 1. Statistical questions with a *p* value of 0.05 were tested via one-way ANOVA (one factor). Two-way ANOVA (two factors) was used to analyse the effect of both deprivation length and repletion type together. Tukey HSD was used to further determine the differences between groups. Statistical analysis and data processing was performed on OriginPro (Version 2021, OriginLab Corporation, Northampton, MA, USA).

## Acknowledgements

This work was funded by UK Research and Innovation (UKRI)’s Biotechnology and Biological Sciences Research Council (BBSRC) grant number BB/N016033/1 and Global Challenges Research Fund (GCRF) as part of the Water Security and Sustainable Development Hub grant number ES/S008179/1. Work in the Saiardi laboratory was supported by the Medical Research Council (MRC) grant MR/T028904/1. The authors thank Dr. Toshikazu Shiba (RegeneTiss, Japan) for the generous supply of polyP100.

For the purpose of Open Access, the author has applied a CC BY public copyright licence to any Author Accepted Manuscript version arising from this submission.

## Author contributions

Experimental design: TZB, AB, AS, MACV; Experimental work: TZB, AS. Data interpretation: TZB, AB, AS, MACV; Funding AB, MACV, AS. Initial draft TZB, AB. All authors contributed to revision and finalisation of the manuscript.

## Data availability

All the data pertaining to this manuscript are in the manuscript or associated supplementary information files.

## Conflict of interest

The authors declare no conflict of interest.

## Supplementary information

**Figure S 1.**
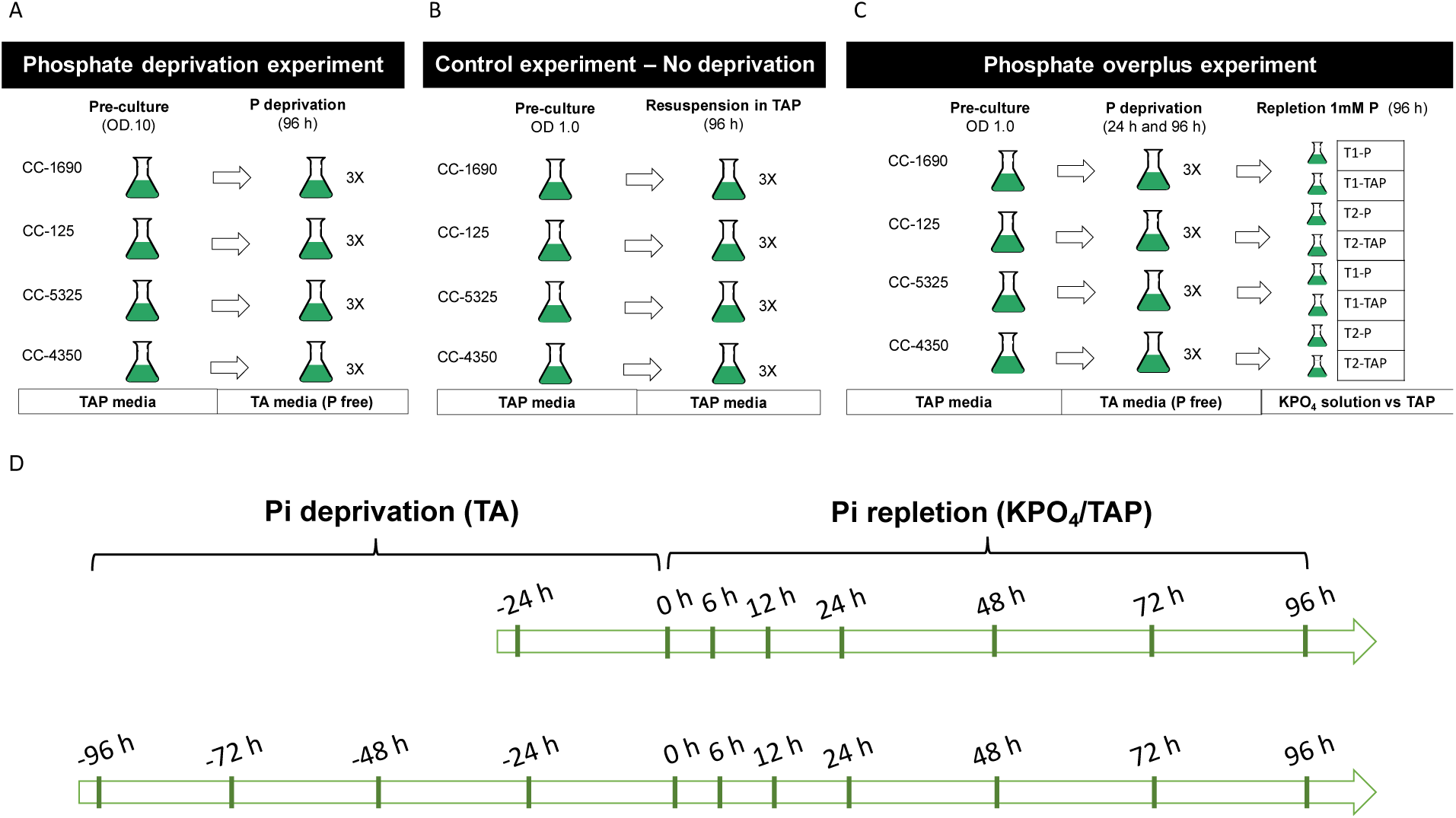
Experimental design diagrams. In all experiments a preculture was grown until an optical density at 750nm of absorbance OD_750nm_= 1.0 was reached. Biomass was harvested (washed) and resuspended in either TAP or TA media. A Phosphate deprivation experiment, Cells were Pi-deprived in TA media for 6 days. B Control experiment. Cells were resuspended in TAP media for 96 h and there was no deprivation of phosphate. C Phosphate overplus experiment. Cells were resuspended in TA media for 24 h or 96 h, then cultures were repleted with either potassium phosphate solution or resuspended in fresh TAP media (both 1mM Pi) for 96 h. D Timeline for sample collection during Pi overplus experiment.

**Figure S2.**
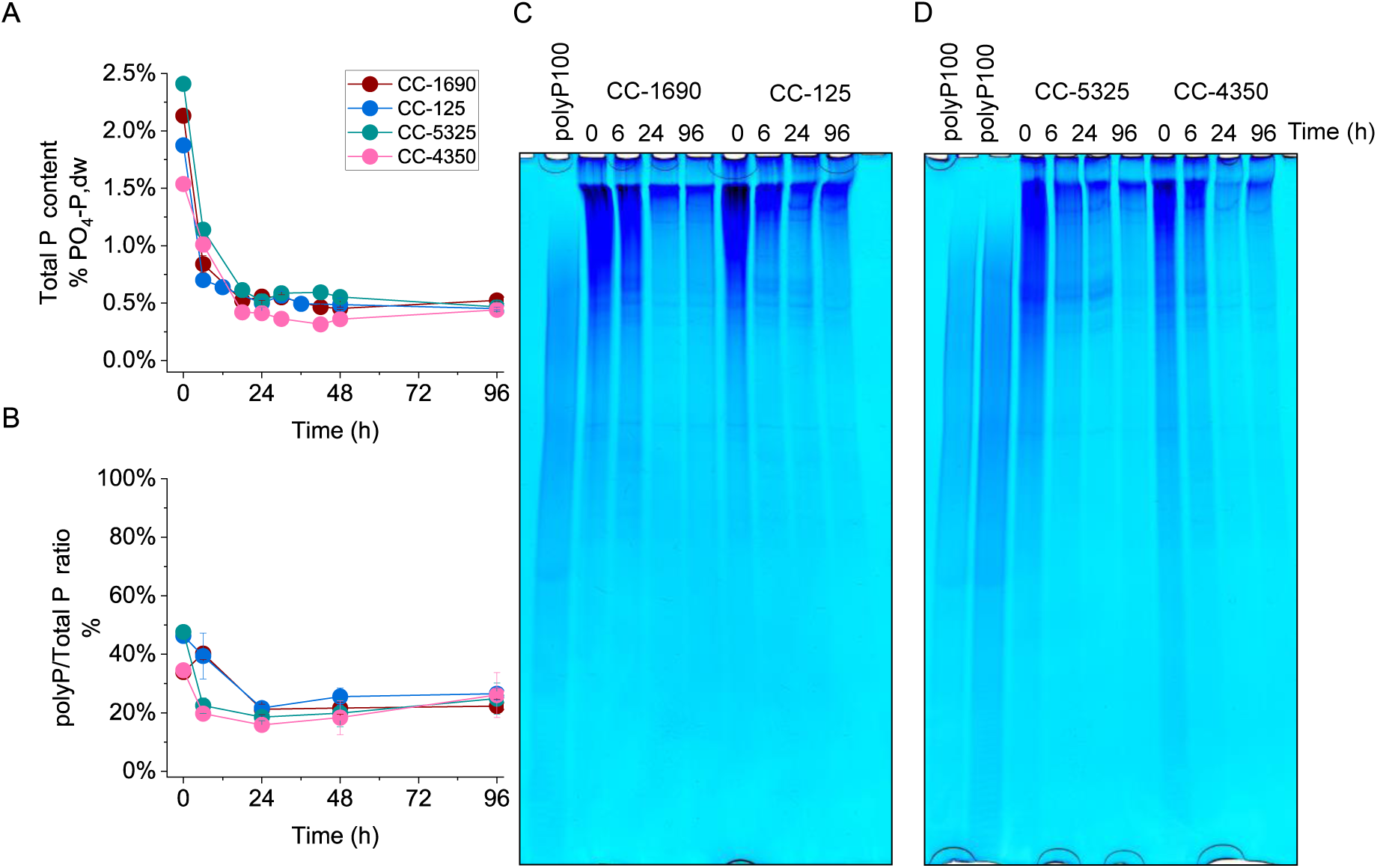
The ratio of polyP to total P falls rapidly during Pi deprivation. A Total P content in biomass (% PO_4_-P,dw), B PolyP to total P mass ratio and C 20% PAGE allows qualitative analysis of polyP. Left, polyP100 loading control, for the four strains of Chlamydomonas the RNA volume equivalent to 500µg dw was loaded on each well for the 0,6, 24 and 96 h time points after OD1.0 cells were harvested and resuspended in TA media.

**Table S 1.** Pi repletion as TAP promotes biomass growth. For the four strains CC-1690, CC-125, CC-5325 and CC-4350 after 24 h and 96 h Pi-deprived cells were harvested and resuspended in fresh TAP media. A Specific growth rates (h^-1^) and B Biomass productivity (g dw L-^1^h^-1^). The data represents the average value (n=3) ± SE.

| A<br>TAP repletion after Pi<br>deprivation<br>Strain | Specific growth rate<br>($\text{h}^{-1}$) | |
| --- | --- | --- |
|  | until min<br>polyP (24 h) | until min<br>RNA (96 h) |
| CC-1690 | $0.034 \pm 0.006$ | $0.024 \pm 0.001$ |
| CC-125 | $0.050 \pm 0.007$ | $0.019 \pm 0.003$ |
| CC-5325 | $0.036 \pm 0.006$ | $0.038 \pm 0.002$ |
| CC-4350 | $0.035 \pm 0.011$ | $0.034 \pm 0.006$ |
| Average | $0.039 \pm 0.004$ | $0.029 \pm 0.005$ |
| B<br>TAP repletion after Pi<br>deprivation<br>Strain | Doubling time<br>(h) |  |
|  | until min<br>polyP (24 h) | until min<br>RNA (96 h) |
| CC-1690 | $39.3 \pm 4.4$ | $33.8 \pm 3.5$ |
| CC-125 | $40.6 \pm 12.2$ | $34.9 \pm 8.3$ |
| CC-5325 | $33.1 \pm 1.2$ | $28.3 \pm 5.8$ |
| CC-4350 | $14.1 \pm 0.9$ | $22.5 \pm 0.9$ |
| Average | $31.8 \pm 6.1$ | $29.9 \pm 2.8$ |
| C<br>TAP repletion after Pi<br>deprivation<br>Strain | Biomass productivity<br>($\text{g dw L}^{-1} \text{h}^{-1}$) | |
|  | until min<br>polyP (24 h) | until min<br>RNA (96 h) |
| CC-1690 | $0.024 \pm 0.002$ | $0.026 \pm 0.001$ |
| CC-125 | $0.039 \pm 0.006$ | $0.020 \pm 0.003$ |
| CC-5325 | $0.021 \pm 0.004$ | $0.030 \pm 0.002$ |
| CC-4350 | $0.021 \pm 0.002$ | $0.023 \pm 0.003$ |
| Average | $0.026 \pm 0.004$ | $0.025 \pm 0.002$ |

**Figure S 3.**
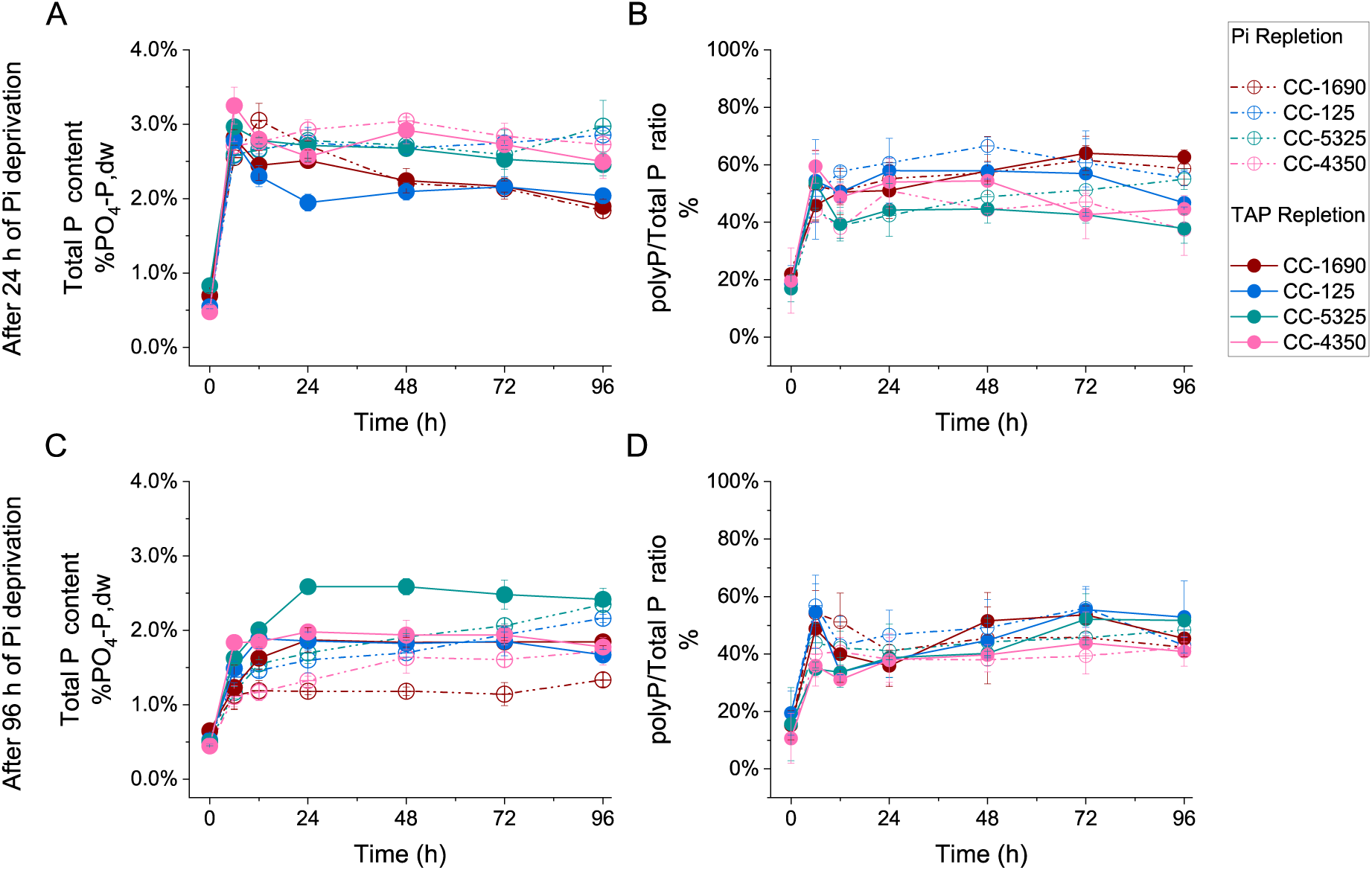
Biomass total P follows similar trend to that of polyP accumulation during Pi overplus. Cells (*C reinhardtii* CC-1690, CC-125, CC-5325 and CC-4350) were grown until OD_750nm_= 1.0, harvested (washed) and resuspended in TA media. After 24 h (A,B) or 96 h (C,D) of Pi deprivation, the cultures were repleted with either KPO_4_ (dashed dotted lines) solution or harvested and resuspended in fresh TAP media (solid lines) (1mM P) and monitored for 96 h. B and D show the corresponding polyP to total P mass ratios (%).

**Figure S 4.**
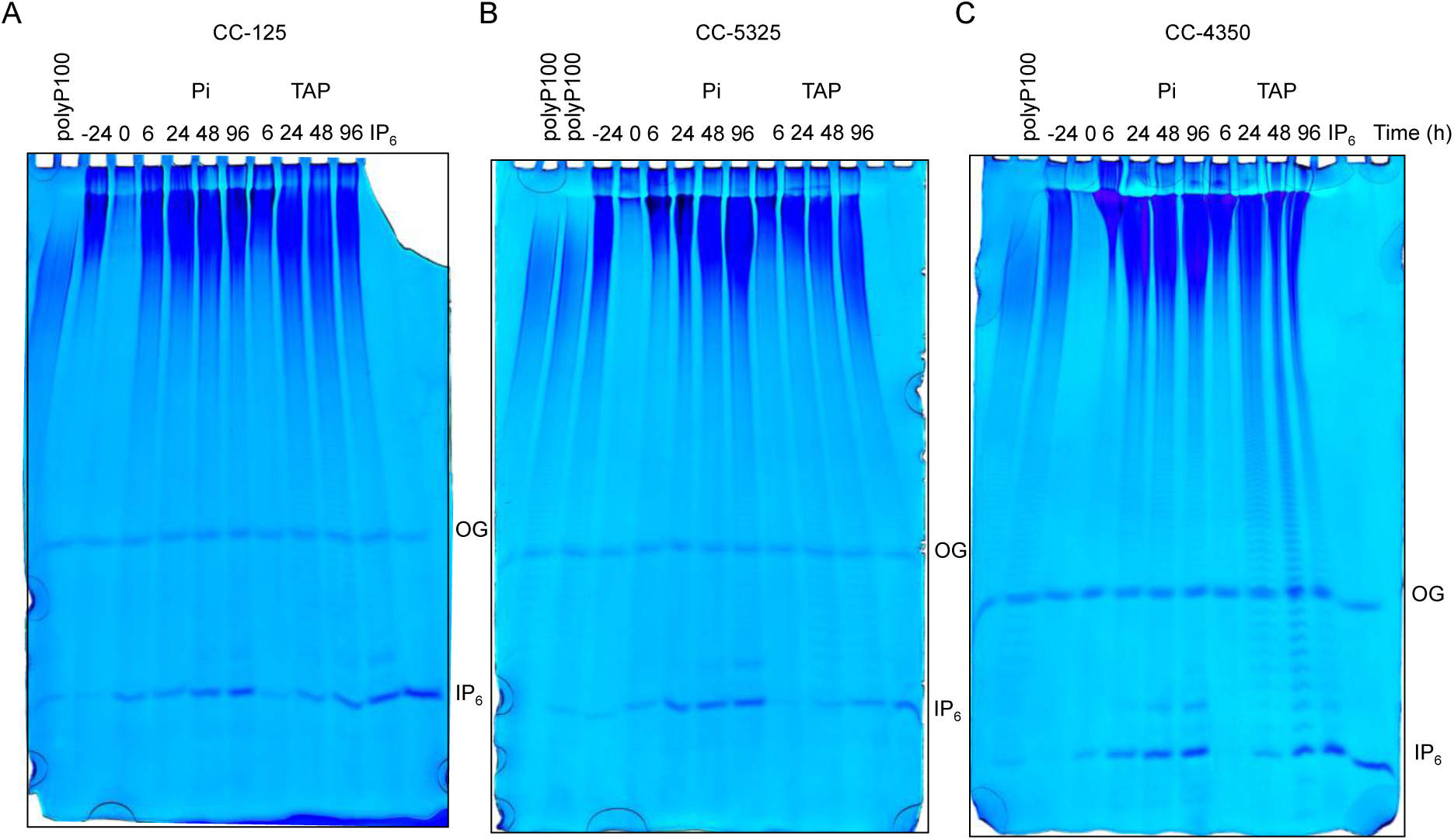
*C. reinhardtii* strains CC-125, CC-5325 and CC-4350 show similar patterns of PolyP and IP_6_ accumulation when repleted with KPO_4_ (Pi) or TAP after 24h Pi deprivation. 33% PAGE. Left, polyP100 loading control, the RNA volume equivalent to 500µg dw was loaded for each sample. Wells from left to right: polyP100 loading control, -24 h (OD_750nm_= 1.0 cells before Pi deprivation), 0 h (24 h Pi-deprived cells before Pi repletion), 6-96 h (After repletion with KPO_4_), 6-96 h (After resuspension in TAP media) and 2 nmol of IP_6_ standard.

**Figure S 5.**
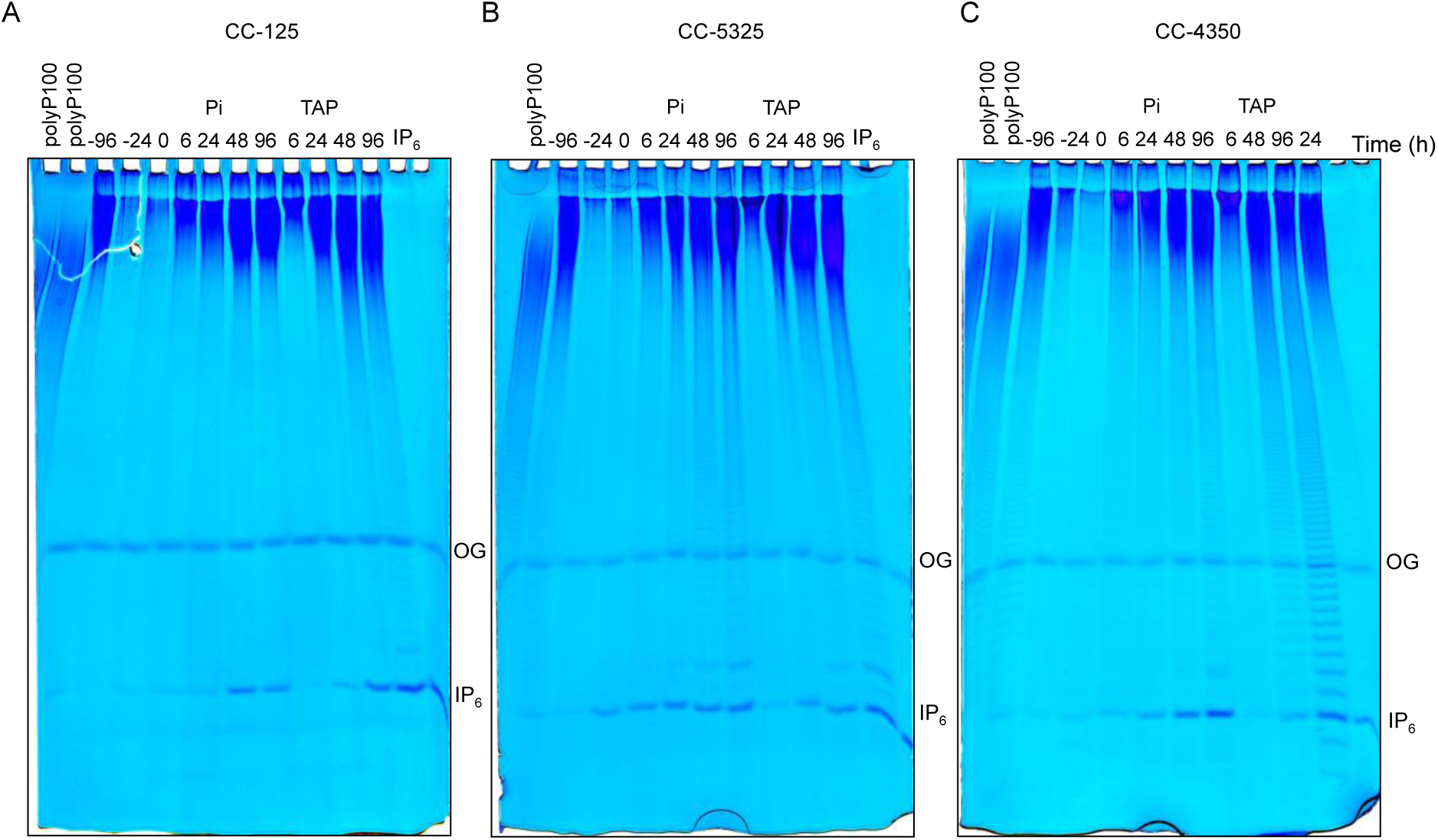
*C. reinhardtii* strains CC-125, CC-5325 and CC-4350 show similar patterns of PolyP and IP_6_ accumulation when repleted with KPO_4_ (Pi) or TAP after 24h Pi deprivation. 33% PAGE. Left, polyP100 loading control, the RNA volume equivalent to 500µg dw was loaded for each sample. Wells from left to right: polyP100 loading control, -96 h (OD_750nm_= 1.0 cells before Pi deprivation), -72 h (After 24 h Pi deprivation), 0 h (96 h Pi-deprived cells before Pi repletion), 6-96 h (After repletion with KPO_4_), 6-96 h (After resuspension in TAP media) and 2 nmol of IP_6_ standard.

**Table S 2.**
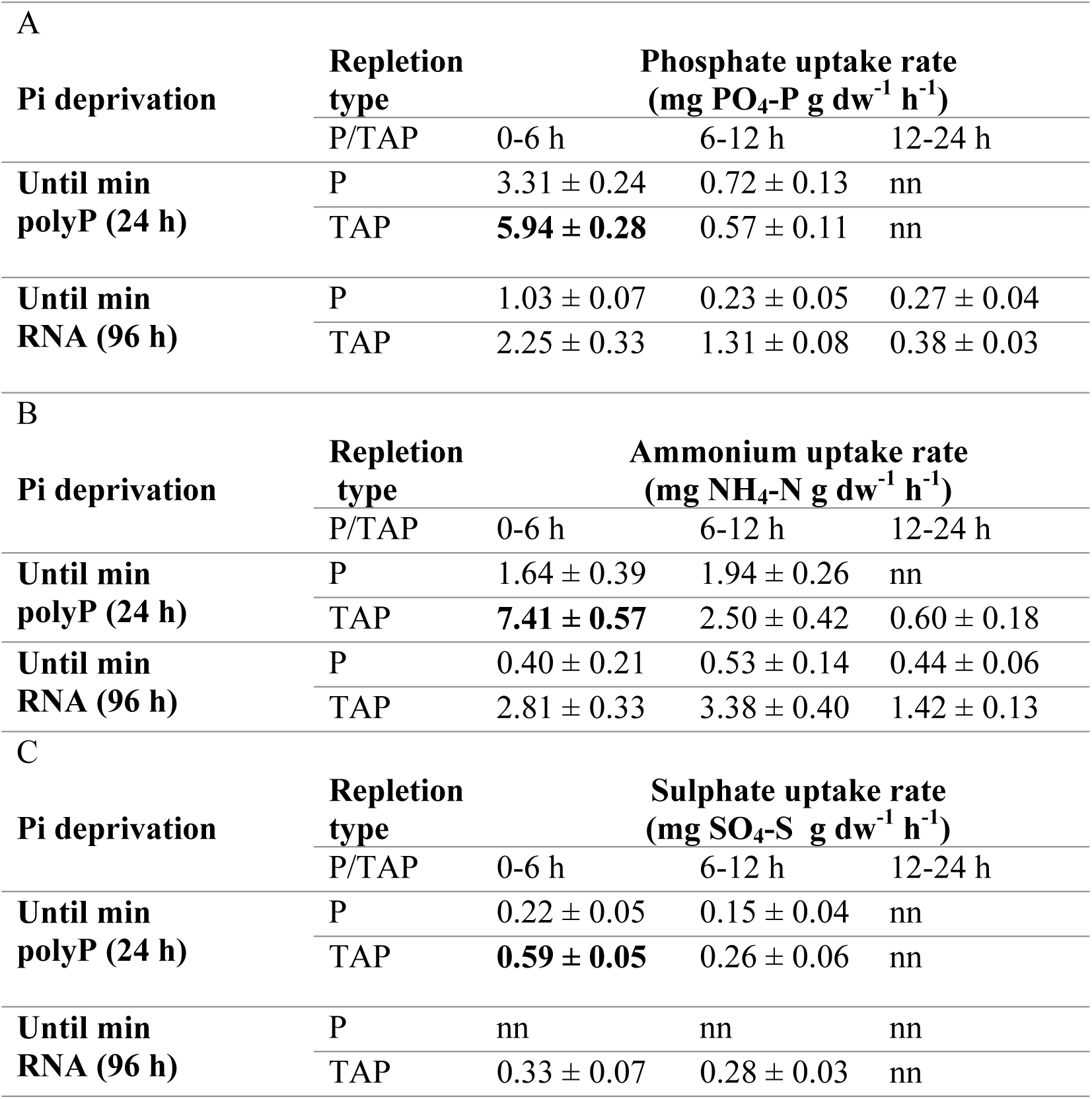
Enhanced nutrient removal was achieved when Pi-deprived cells were repleted with complete nutrients. For both periods of Pi deprivation (24 and 96 h) and KPO_4_ (Pi) vs TAP repletion, the average value of the four strains of *C. reinhardtii* (CC-1690, CC-125, CC-5325 and CC-4350) was used, and the nutrient uptake rates of phosphate (PO_4_-P), ammonium (NH_4_-N) and Sulphate (SO_4_-S) were calculated. The nutrient uptake rates are shown for the first 24 h of phosphate repletion (6, 12 and 24 h). The ‘nn’ values represent the nutrient uptake rates lower than 0.1 mg g dw^-1^ h^-1^.

**Figure S 6.**
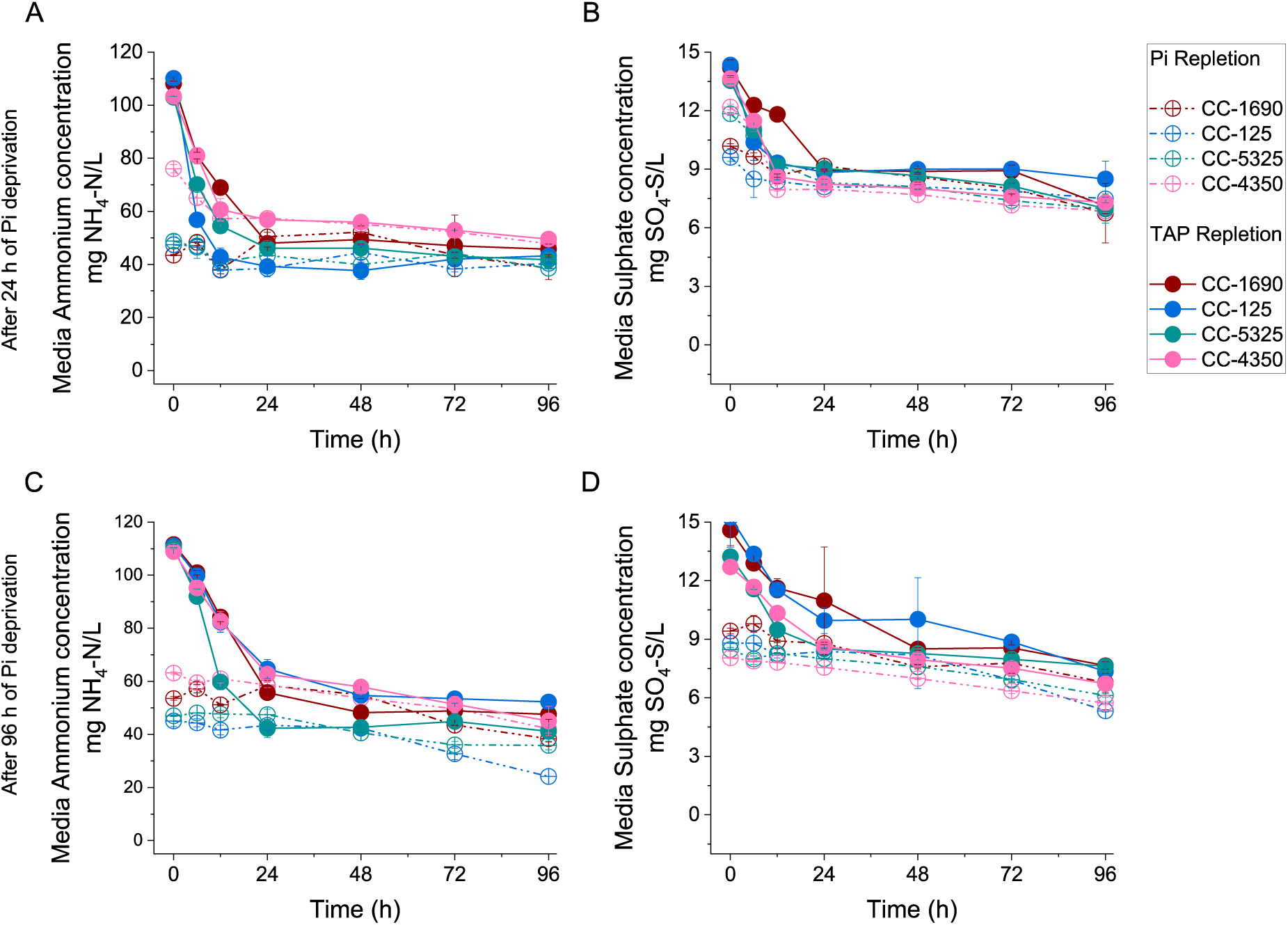
Ammonium and Sulphate removal from the media during Pi overplus. Cells of *C. reinhardtii* (CC-1690, CC-125, CC5325 and CC-4350) were grown until OD_750nm_= 1.0, harvested (washed) and resuspended in TA media. After A and B 24 h or C and D 96 h of Pi deprivation, the cultures were repleted with either KPO_4_ (dashed dotted lines) solution or harvested and resuspended in fresh TAP media (solid lines) (1mM Pi) and monitored for 96 h. A and C Ammonium concentration in the media (mg NH_4_-N/L). B and D Sulphate concentration in the media (mg SO_4_-S/L).

